# Auditory cortex deviance detection represents a canonical difference signal

**DOI:** 10.64898/2026.01.06.697993

**Authors:** Adam Hockley, Connor G Gallimore, Jordan P Hamm, Manuel S Malmierca

**Affiliations:** Cognitive and Auditory Neuroscience Laboratory, Institute of Neuroscience of Castilla y León (INCYL), Salamanca, Spain 37007; Salamanca Institute for Biomedical Research (IBSAL), Salamanca, Spain 37007; Department of Cell Biology and Pathology, Faculty of Medicine, University of Salamanca, Salamanca, Spain 37007; Emotional Brain Institute, Nathan S. Kline Institute for Psychiatric Research, Orangeburg, NY, USA 10962; New York University Grossman School of Medicine, New York, NY, USA 10016; School of Psychology and Neuroscience, University of Bristol, Bristol, UK BS8 1TD

**Keywords:** Predictive coding, prediction error, deviance detection, context, auditory cortex, oddball

## Abstract

Context modulates neural processing of sensory stimuli. Excitatory neuron responses are most often suppressed to stimuli that are typical of the current context and augmented to stimuli that are deviant. These effects have been extensively studied in basic and clinical neuroscience using oddball paradigms, involving the repeating of standard stimuli and the insertion of rare deviant stimuli. Augmented neural responses to deviants - referred to as deviance detection - have been interpreted within predictive coding frameworks as prediction errors, signaling the difference between internal predictions and actual sensory input. However, most oddball paradigms simultaneously alter both predictions and input, so cannot test for a true difference signal. As yet, no studies have demonstrated that neural deviance detection during oddball paradigms specifically encodes the difference between internal predictions and unexpected sensory input, as opposed to, e.g., a generalised surprise signal or augmented bottom-up signaling. Here we compare neural responses to auditory stimuli across oddball paradigm variants. In a classical oddball paradigm, responses of putative excitatory neurons in the primary auditory cortex (A1) to auditory deviants contain information about recent frequency change and retain a memory trace of previous stimulus context. Interestingly, in a fixed-deviant oddball paradigm where the standards (theoretical predictions) were altered between blocks, but deviant stimulus features remain unchanged, neural response patterns encoded standard-to-deviant frequency difference. Together, these findings support the interpretation that A1 deviance detection contains a sensory prediction error that represents the difference between prediction and sensory input, a corollary of the predictive coding framework.

## INTRODUCTION

Sensory context modulates sensory processing in the mammalian brain. This modulation has been studied across sensory modalities using “oddball” paradigms consisting of repeating standard stimuli (STD; contextually redundant) and unpredictable deviants (DEV; contextually deviant). Augmented neural responses to deviant stimuli if known as “deviance detection”, and is present at the single neuron level as early as primary sensory cortex (Dürschmid et al., 2016; Parras et al., 2017; Audette et al., 2022; Gelens et al., 2024; Hockley et al., 2025; Shymkiv et al., 2025), but it has most often been quantified as “mismatch negativity” (MMN) in human EEG studies, where it is considered a reliable biomarker of cognitive function and dysfunction in several neuropsychiatric disorders such as schizophrenia and autism spectrum disorders. Basic studies of neural deviance detection, in relation to other context-based modulations present during the oddball (Fig 1A), are therefore paramount for advancing understanding of neural pathology in neuropsychiatry.

**Figure 1:**
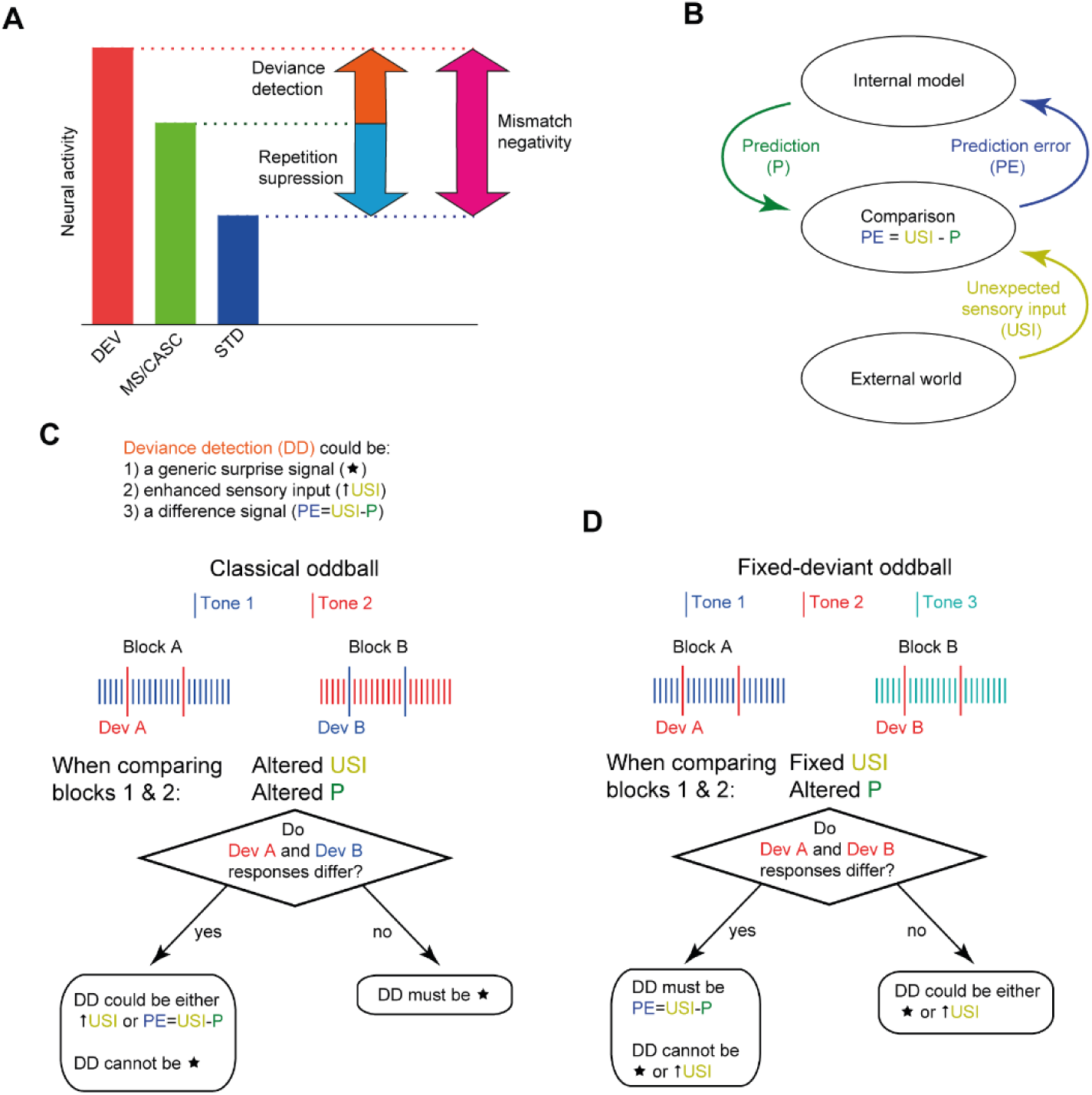
A) A schematic for how the strength of neural responses to deviants (DEV), controls such as many standard or cascade (MS/CASC), and standards (STD) are widely used to calculate deviance detection (DD = DEV - MS/CASC), repetition suppression (RS = MS/CASC - STD) and mismatch negativity (MMN = DEV - STD). **B)** The classical predictive coding model. **C)** How classical oddball paradigms can reveal the information contained by cortical deviance detection, when understood under the predictive coding framework. Deviance detection could putatively convey generalised surprise (◻; i.e. a generic neural response when anything unexpected occurs), an enhancement of the unexpected sensory input (◻USI), or a difference between input and internal predictions (PE=USI-P). The classical oddball we employ alters components of the predictive coding framework, when comparing between blocks. Comparing deviants in the classical oddball theoretically studies altered predictions and unexpected sensory input. **D)** As in C; how fixed-deviant oddball paradigms can reveal the information contained by cortical deviance detection, when understood under the predictive coding framework. Comparing deviants in the fixed-deviant oddball theoretically alters predictions, while unexpected sensory input remains constant. A combination approach of classical and fixed-deviant oddballs affords us to determine information conveyed by deviants.

Predictive coding has been proposed as a theoretical framework to explan sensory contextual processing during oddball paradigms, where the brain encodes regular patterns, generates an internal predictive model, and then detects deviations from the resulting predictions (Rao and Ballard, 1999; Friston, 2005; Clark, 2013; Keller and Mrsic-Flogel, 2018). However, experimentally testing the neural mechanisms underlying this framework remains an active area of research (Aizenbud et al., 2025). Within this framework, neural deviance detection (and human MMN) is interpreted as a “sensory prediction error” (Dürschmid et al., 2016; Parras et al., 2017; Audette et al., 2022; Gelens et al., 2024; Hockley et al., 2025; Shymkiv et al., 2025): a theoretical signal produced when an external input does not match internal predictions, potentially triggering the updating of internal models (Fig 1B). The neural mechanisms of deviance detection include hierarchically organised circuitry, both within cortical layers (Arnal and Giraud, 2012; Bastos et al., 2012; Gallimore et al., 2023) and between brain regions (Hamm et al., 2021; Bastos et al., 2023; Asko et al., 2024; Han and Helmchen, 2024; Hockley et al., 2025). This hierarchical organisation is consistent with predictive coding theories that posit top-down predictive modulation and bottom-up prediction error signals. Classic frameworks also present prediction errors as a difference between an unexpected input and internal predictions (Rao and Ballard, 1999; Friston, 2005; Clark, 2013). However, while evidence of a difference calculation has been shown in reward prediction errors (Schultz, 2016; Watabe-Uchida et al., 2017; Gershman et al., 2024), direct evidence for an equivalent computation underlying sensory prediction errors remains elusive.

The notion that sensory prediction errors signal differences between actual inputs and internal predictions has been tested (and not found) in the mouse visual cortex in relation to spatial context and visual deviants, during navigation (Muzzu and Saleem, 2021; Furutachi et al., 2024). Whether deviance detection during conventional auditory oddball paradigms concord with predictive coding theories in this manner – i.e., whether they reflect differences between current and expected inputs – remains to be rigorously tested at the single neuron level, particularly within the auditory cortex.

What information neural deviance detection passes upstream within the brain, particularly in sensory context processing paradigms like the oddball, is not wholly understood (for review, see (Furutachi and Hofer, 2026)). Addressing this unanswered question is essential to understanding whether deviance detection arises from predictive coding-like functions in hierarchical brain circuits. Moreover, identifying precisely what information is encoded by deviance detection would provide a critical opportunity to test biological accuracy and neural mechanisms of classical predictive coding theories.

Here we interrogate a foundational assumption of the predictive coding framework: that neural deviance detection is computed as a difference between sensory input and internal predictions. To achieve this, we study the information encoded by A1 neurons during classical and fixed-deviant auditory oddball paradigms which alter theoretical predictions and prediction errors, between blocks. We examine information encoded through both temporal firing patterns at the single neuron level, and spatial patterns of neural activity at the population level. We first use a classical oddball (Fig 1C) to reveal that deviance detection is not a generalised surprise signal, but carries stimulus-relevant information (Fig 2). Next, using a cascade paradigm we show that individual A1 neurons strongly contain changes in sound frequency, while only weakly representing absolute frequency information (Fig 3). Finally, we use a fixed-deviant paradigm which theoretically alters predictions while maintaining sensory input constant (Fig 1D). This approach rules out that deviance detection conveys enhanced sensory information or generalised surprise, instead revealing that neural deviance detection contains information about the difference between predicted and actual sensory input (Fig 4). Together, these findings provide neural evidence that auditory deviance detection contains a foundational hallmark of classical predictive coding frameworks: information about the mismatch between sensory inputs and internal predictions.

**Figure 2:**
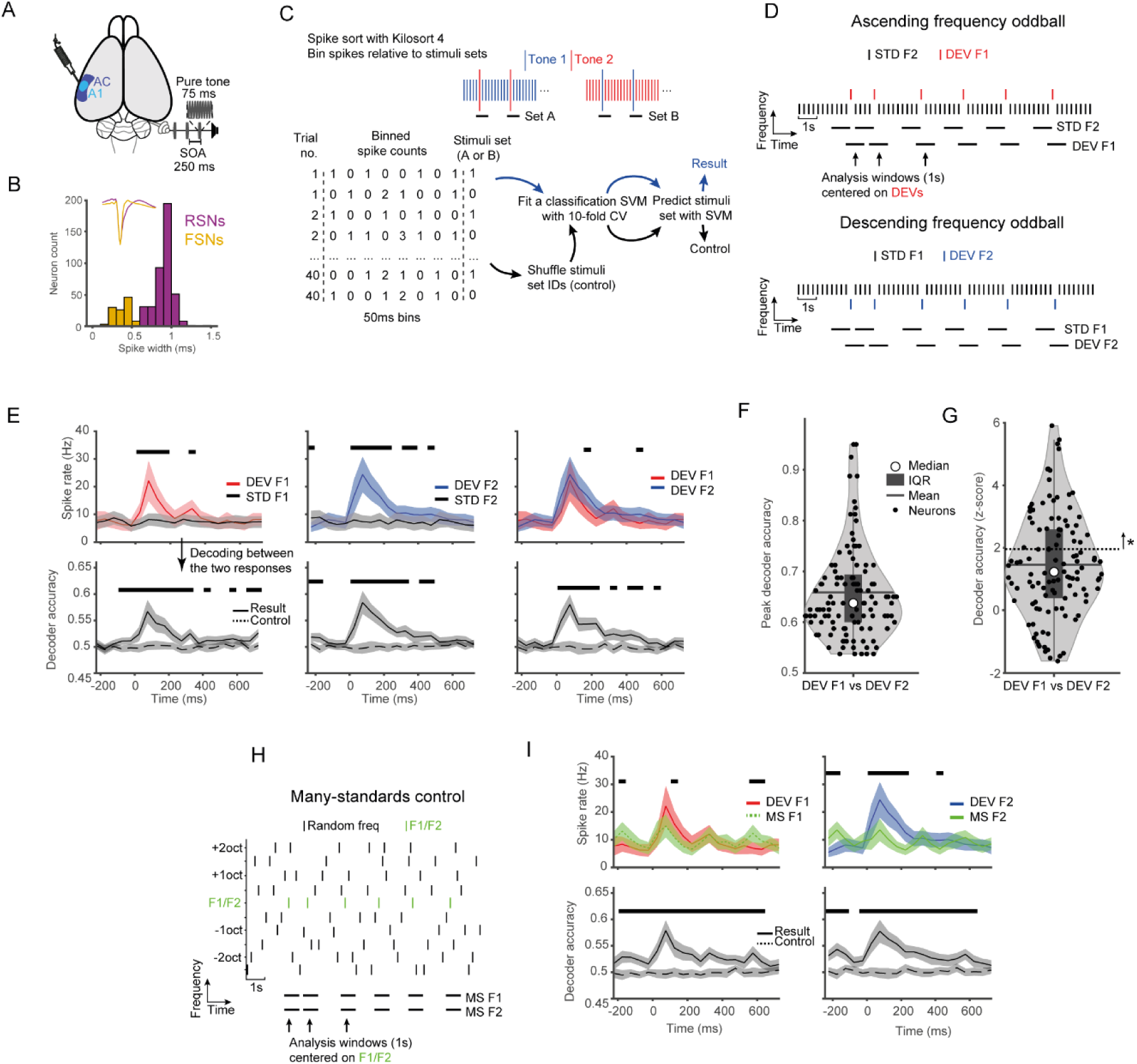
Deviance detection does not signal a generalised surprise signal. **A)** Schematic of multichannel recording. **B)** A1 neurons were separated by spike width (positive to negative peak duration) into regular spiking (RSNs; putative excitatory) and narrow spiking (FSNs; putative inhibitory) neurons. **C)** SVM decoder methodology. **D)** Data compared were 40 analysis windows containing the same stimuli within each stimulus block. During a frequency auditory oddball paradigm, two different frequencies are presented (F1 & F2). We compared stimuli of 4 repeated standards (STD F1 & STD F2) to deviants (DEV F1 & DEV F2) to assess context encoding. **E)** Top: PSTH responses. Bottom: Decoder accuracy classifying between the two responses above. Shaded areas represent 95% CI. **F)** Violin plot of peak decoding accuracy between F1 and F2 deviants. **G)** Violin plot of the decoding between F1 and F2 deviants. Dotted line is z-score 1.96. **H&I)** Stimuli from the many-standards control, centered on F1 or F2 were compared to deviants using the same methodology as above. Black bars indicate p < 0.01, paired t-test. n = 116 mismatch RSNs.

**Figure 3:**
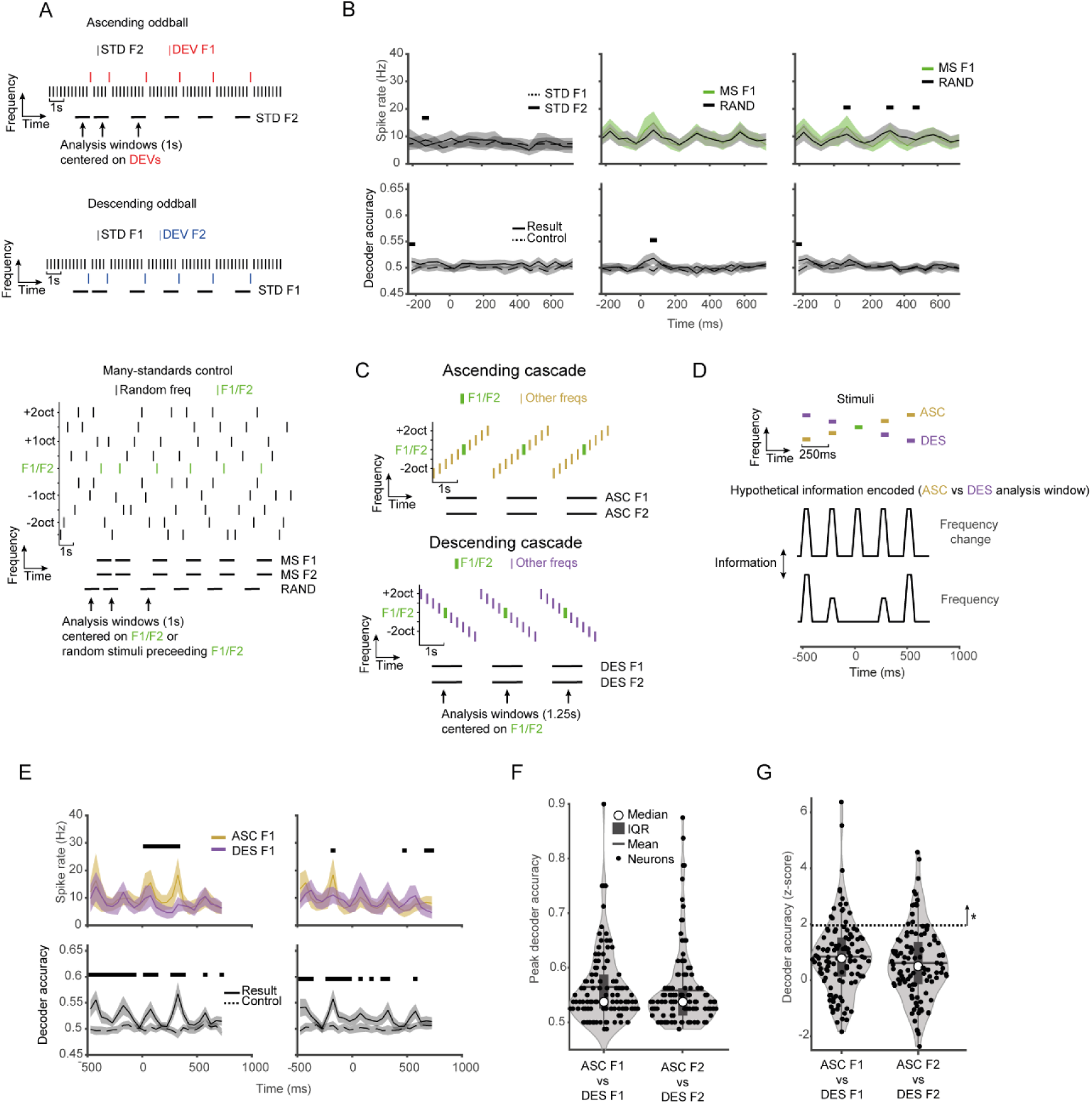
Encoding of frequency change information. **A)** We compared stimuli to reveal encoding of frequency information. Analysis windows consisting of 4 repeated STD tones were compared for F1 & F2. For the many-standards control, analysis windows centered on F1 or F2 were compared to windows containing random frequencies. **B)** Top: PSTH responses. Bottom: Decoder accuracy classifying between the two responses above. **C)** Analysis windows were centered on each frequency (F1 or F2) within ascending and descending cascades. **D)** Schematic of hypothetical frequency and frequency change information when comparing ascending and descending cascades. **E)** Top: PSTH response. Bottom: Decoder accuracy classifying between the two responses above. Shaded areas represent 95% CI. Black bars indicate p < 0.01, binwise paired t-test. **F)** Violin plot of peak decoding accuracy between ascending and descending cascades (0-200 ms time window). **G)** Violin plot of decoding between ascending and descending cascades. Dotted line is z-score 1.96. n = 116 mismatch-responsive RSNs.

**Figure 4:**
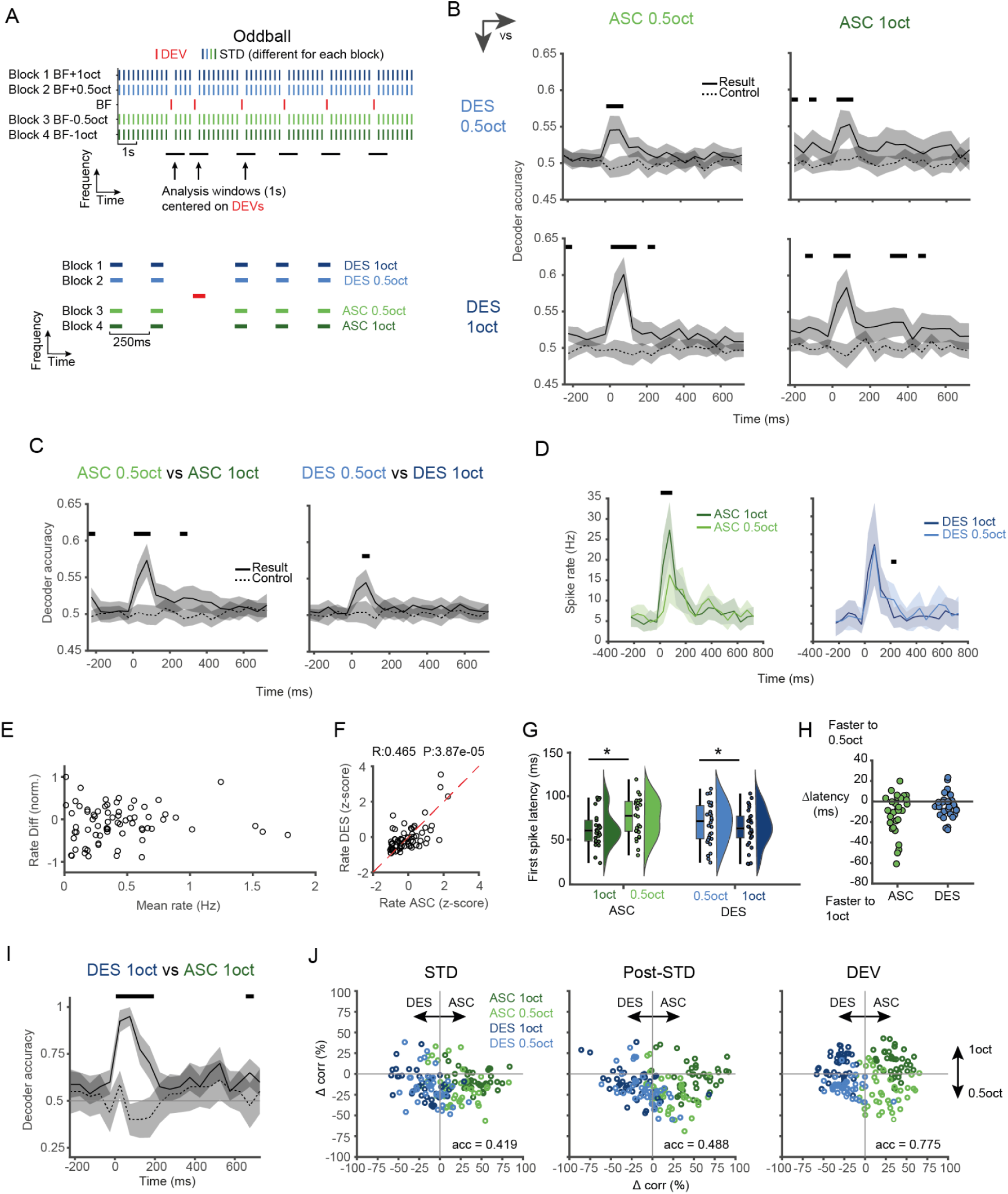
A1 RSN deviance detection encodes the difference between prediction and sensory input. **A)** Oddball stimuli of a fixed frequency deviant and variable standards. **B)** Grid comparisons of ascending vs descending oddballs. Shaded areas represent 95% CI. Black bars indicate p < 0.01, binwise paired t-test. **C)** Comparisons for ascending or descending oddballs of different frequency steps. **D)** Spike rate data for deviant responses during varied standard stimuli. **E)** Scatter plot for mean rate (ascending and descending 1 octave DEVs) vs rate difference (ascending – descending; normalised). **F)** Scatterplot of z-scored evoked rates by ascending and descending 1 octave deviants. **G)** Raincloud plots of first spike latencies for the 4 different conditions. **H)** Latency differences for individual neurons. **I)** Population decoding deviant responses between individual trials of ASC 1oct and DES 1oct. n = 40 trials. **J)** Population decoding deviant responses between individual trials into average patterns of full/half octave or ASC/DEC. n = 40 trials per deviant type.

## MATERIALS AND METHODS

### EXPERIMENTAL MODEL AND STUDY PARTICIPANT DETAILS

All experiments were conducted at the Cognitive and Auditory Neuroscience Laboratory, Institute of Neuroscience of Castilla y León (INCYL), Salamanca, Spain and performed animal were experiments under protocols established by the Directive of the European Communities (86/609/CEE, 2003/65/CE, and 2010/63/UE) and RD53/2013 Spanish Legislation for the use and care of animals, and this study was approved by the Bioethics Committee of the University of Salamanca (USAL-ID-887). 8 female rats (Long-Evans; 200-280g) from Janvier Labs (Le Genest-Saint-Isle, France) were housed on a 12/12 h light-dark cycle, with food and water freely available. Raw data from 6 of these rats were previously published with entirely different analyses (Hockley et al., 2025).

### METHOD DETAILS

#### Surgery

Neural recording experiments were conducted inside an acoustically insulated and electrically protected room. Anesthesia was induced and maintained with urethane (1.9 g kg^−1^, i.p.; Sigma-Aldrich, St Louis, U.S.A.) with supplementary doses (∼0.5 g kg^−1^, i.p.) administered based on corneal and rear paw withdrawal reflex. Animals were artificially ventilated with O_2_ and temperature was maintained at 36 - 38°C using a rectal probe and a homeostatic heating blanket (Cibertec, Madrid, Spain). Rats were continuously monitored throughout the experiment and periodically administered isotonic glucosaline solution to avoid dehydration (5 ml every ∼8 hours, s.c.). Initially, auditory brainstem responses (ABRs) were recorded by placing subdermal electrodes at the dorsal midline close to the neural crest and behind each pinna. ABRs stimuli were tone bursts (4, 8, 16 and 32 kHz; 5 ms duration, 1 ms rise/fall times, 21 Hz presentation rate, 512 repetitions in 10 dB steps from 0–80 dB SPL; Tucker-Davis Technologies RZ6 & Medusa 4Z preamp). ABR thresholds were visually confirmed as normal hearing (≤40 dB SPL) in all animals.

Following ABRs, dexamethasone was administered (Cortexonavet, 0.25 mg kg^−1^, i.m.; Syva, León, Spain) and atropine (0.1 mg kg^−1^, s.c.; Braun, Rubi, Spain) to reduce cerebral oedema and bronchial secretions, respectively. Pinna cartilage was removed to obtain easier access to the ear canal and performed a tracheotomy to allow artificial ventilation. The animal was then placed in the stereotactic frame with a bite bar and hollow ear bars into the ear canals. The speaker was inserted into the right ear bar and filled the hollow left ear bar with plasticine. A midline incision exposed the skull, the temporalis muscle was removed for A1 access, and a craniotomy allowed access to A1. The dura mater was removed and kept the exposed brain surface warm and moist by application of agar. The posterior fossa was opened to reduce respiratory movements of the brain.

#### Neural recordings

Stimulus patterns were generated and presentation controlled via TDT Synapse software and TDT RZ6 hardware using custom MATLAB (Mathworks, Natick, U.S.A) code. Multi-channel recording probes (H3, Cambridge NeuroTech, Cambridge, UK) were aimed stereotaxically and advanced slowly (see (Fiáth et al., 2019)) through the craniotomy into the target brain area using an IVM Single Microdrive (Scientifica, Uckfield, UK). Signals were amplified by a TDT PZ5 preamp connected to a TDT RZ2 processor, filtered (low-pass 7 kHz), and stream data was recorded using TDT Synapse software at a sampling rate of 24 kHz. For A1 recordings, A1 location was identified using tonotopy from a map developed previously (Nieto-Diego and Malmierca, 2016) and advanced electrodes manually to the pia mater surface then advanced 1300 µm using the Microdrive at 3 µm/s to position all electrode sites within the cortical layers.

#### Stimuli

Sound stimuli were controlled by Tucker-Davis Technologies (TDT; Alachua, FL, USA) hardware and presented unilaterally to the right ear through a closed system using a custom-made earphone (calibrated to ±2 dB at 0.5 - 44 kHz) coupled to a custom-made cone ear bar.

First, receptive-field stimuli consisting of tone bursts were presented (75 ms duration, 250 ms stimulus onset asynchrony, 1−44 kHz, 1/6 octave spacing, 0-70 dB SPL, 10 dB steps, 5 repeats, randomised). From these, a frequency-response area was plotted, and a frequency pair of two tones (F1 & F2; 1/2 octave spaced), and intensity (50-70 dB SPL; 20-40 dB above unit thresholds) was automatically chosen that was centered around the average BF of simultaneously recorded neurons and thus could activate all neurons on the electrode approximately equally. The F1 & F2 frequency pair was used for a classical auditory oddball paradigm where both F1 and F2 were presented as either the standard or deviant in different recording blocks (Fig. 1C; 40 deviants ascending, 40 deviants descending, 10% deviant probability, minimum of 4 standards before deviant).

To assess whether responses to deviant stimuli are true deviance detection and not due to adaptation, a many-standards control was used (Schröger and Wolff, 1996; Ruhnau et al., 2012; Harms et al., 2014), consisting of 10 tones of equally spaced tone frequencies, containing F1 and F2 stimuli, presented in an unpredictable pseudorandom order where a single tone was never repeated (Fig 2H). The many-standards control was used to assess whether the difference between deviant and standard responses reflected predictive activity and/or stimulus specific adaptation. In the many-standards control, each tone has the same low probability (10%) as a deviant in the oddball paradigm, so is unpredictable and unrepetitive, and therefore free of repetition suppression and prediction errors (Parras et al., 2017). Therefore, if the observed difference between deviant and standard responses is caused entirely by repetition suppression to the standard tone, responses to deviant and many-standards controls would be equal given their unrepeated, fixed, low likelihood of occurrence. Inversely, if the observed difference between deviant and standard responses is produced entirely by prediction error, responses to standard and many-standards controls would be equal, with increased deviant responses from prediction error. Our many-standards control used 10 half-octave spaced frequencies from the receptive-field stimuli, centered around the frequency pair (F1 & F2, which would be stimuli 4-5 or 5-6 depending on their frequency, to allow the many standards stimuli to remain within the hearing range of the rat) presented in a pseudorandom order with no stimulus repetition. Cascade stimuli consist of a sequence of the same 10 tones, ascending or descending in frequency order and are composed of frequency and change direction information. The F1 and F2 stimuli were included in the many-standards and cascade controls.

In the second experiment, receptive fields were calculated as above. A fixed-deviant oddball paradigm with an invariant deviant and varied-frequency standard stimuli was used (Fig 1D). This is optimised for testing a difference signal, as it theoretically alters predictions while maintaining sensory input constant (Furutachi et al., 2024). Our auditory fixed-deviant oddball paradigm used a single deviant at the average best frequency of all neurons in the recording, alongside 4 different standards that were used in separate recording blocks, at +1, +0.5, -0.5 and -1 octaves from the deviant frequency (Fig. 4A; 40 deviants for each condition, 10% deviant probability, minimum of 4 standards before deviant).

### EXPERIMENTAL DESIGN AND STATISTICAL ANALYSES

#### Neural recordings analysis

TDT stream data were analyzed as single units using a custom analysis pipeline (https://github.com/adam-hockley/TDT_Single_Unit_Pipeline). This combined the streams from multiple TDT recording blocks i.e. all stimuli presented for one electrode position, into a single binary file. The binary data stream was then processed in SpikeInterface (Buccino et al., 2020), involving preprocessing (0.3-3 kHz filter, common median reference), and spike-sorting with Kilosort4 (Pachitariu et al., 2024). The pipeline then converted output data in Phy format back into a TDT block-like format for further analyses relative to the timings of auditory stimuli presented. Only ‘good’ units from the sorter output were used for these analyses.

Coding of contextual deviance was assessed across the layers of A1 by separating all neurons into three subdivisions based on depth from cortical surface: supragranular (SG; 0-300µm), granular (G; 320-600 µm) and infragranular (IG; 620-1280 µm). These depths putatively represent layers I, II, and the top half of layer III (SG), bottom half of layer III and layer IV (G), and layers V a VI (IG), and were based on histological layer identification (Smith et al., 2012).

For the classical oddball data, mismatch-responsive regular spiking neurons (RSNs) were identified as neurons that could differentiate between standard and deviant responses, when compared to a shuffled condition that was repeated 100 times. Standards used were the only final standard before a deviant, to equate for the number of trials across conditions. The decoder accuracy z-score was computed in reference to the shuffled condition, and significant mismatch encoding for individual neurons was confirmed if the z-score was >1,96 for either F1 or F2 stimuli (>95% CI). For the fixed-deviant data, mismatch responsiveness was also determined by significant decoding between standard and deviant responses compared to a shuffled condition. Significant mismatch encoding for individual neurons was confirmed if the z-score was >1,96 for either descending or ascending 1 octave conditions.

To test the number of neurons that were significantly differentiating between ascending and descending deviants, the 0-100ms time window was further analyzed. The same decoder was then compared to a shuffled condition that was repeated 100 times. The decoder accuracy z-score was computed in reference to the shuffled condition, and significant encoding for individual neurons was confirmed if the z-score was >1.96 (the 95% CI for standard distribution).

Analysis of first spike latency (FSL) was restricted to only mismatch responsive, broad-spiking single units. An evoked response was defined as occurring between 10 and 150ms of stimulus onset, and only neurons which exhibited a spike within this window in at least 10 trials were included to ensure a robust mean FSL was calculated for each neuron. Thirty mismatch-responsive single units met this criteria, and their average latency was estimated by taking the median first spike latency across all trials within each condition (ASC-OCT, ASC-hOCT, DES-hOCT, DES-OCT). Statistical significance was assessed using a two-way ANOVA with deviant frequency magnitude (hOCT or OCT) and direction (ASC or DES) as within-subjects factors. Significant main effects were followed up with simple effects analysis using paired sample t-tests.

#### Decoding and statistical analysis

Decoding analyses were used to determine when evoked responses of single neurons or populations of neurons differentiated between different stimuli or the same stimuli in different contexts. Binwise decoding of stimulus information was used individually for each neuron. Spike timing data were divided into datasets with timings relative to certain stimuli. Spike times were time-binned (Matlab function histcounts; bin width 50ms), and used as the predictors for the neuronwise binwise classification SVM, with the stimuli as the response values. The 50ms bin size is too small to detect precise spike timing differences, but the auditory cortex largely relies on a rate code (Buck et al., 2025) and frequency is encoded by rates in A1 (Micheyl et al., 2013). For each bin a classification SVM was fit to compare spike times from two different analysis window datasets (Matlab function fitcsvm; Radial basis function kernel; standardised; 10-fold cross-validation). Decoder accuracy was calculated on data not used for training (Matlab function kfoldPredict). In a control condition, stimulus identifiers were shuffled before fitting the SVM classifier.

For the fixed-deviant data in Fig 4 I-J, population decoding was conducted using a trial-wise Maximum correlation classifier, where for all neurons in the dataset, two mean patterns were calculated based on either deviant type (i.e. DES 1oct & ASC 1 oct), or Magnitude (full vs half) or change Direction (ASC vs DES). Population data from individual trials were compared to mean patterns and the trial was assigned to either of the patterns based on that with the highest correlation coefficient. This analysis was conducted for individual bins, allowing comparison of STD (-250 to -200 ms) post-STD but pre-DEV (-100 to -50 ms) and DEV (0-50 ms) time windows. Accuracy scores when decoding Magnitude and Direction were calculated as the proportion of trials classified correctly for both decoders (chance = 0.25).

Plotting and statistical tests for decoder accuracy were performed using custom publicly available MATLAB scripts. Binwise two-sample t-tests (α = 0.01) and linear mixed-effect models (α = 0.05) were used to compare between single-neuron decoder accuracy for the normal and shuffled control conditions. First spike latencies (FSLs) were compared using two-way repeated-measures ANOVAs. Binwise two-sample t-tests (α = 0.01) were used to compare population decoder accuracies between normal and shuffled control conditions.

### CODE ACCESSIBILITY

All data are available at 10.5281/zenodo.18166301. All analysis code is available at 10.5281/zenodo.17516163. Any additional information required to reanalyze the data reported in this paper is available from the corresponding author upon request.

## RESULTS

### A1 neurons encode contextual deviance

We aimed to compare how neurons in A1 encode information about sound frequency and context during the auditory oddball paradigm. We recorded single units from A1 (n = 637 units total from 35 electrode positions; 2240 total channels; n = 6, female rats). First, we mapped receptive fields, then designed a classical oddball paradigm for each electrode location using two half-octave spaced tones (F1 & F2), both equal intensity, suprathreshold, within the receptive field and thus could activate all neurons on the electrode approximately equally (see methods for receptive field frequencies and F1/F2 frequency picking). F1 and F2 varied in frequency depending on the tonotopy of the recording location. In the classical oddball, we recorded two blocks where we used each tone (F1 and F2) as the standard in one block and as the deviant in another (i.e., a flip-flop) to allow comparison of the chosen tones for either context (standard or deviant). F1 and F2 varied in frequency depending on the tonotopy of the recording location. Oddball sequences were either ascending or descending, depending on whether the deviant was a higher or lower frequency than the standard, respectively (Fig. 2D). We then a) compared mean spike rate responses between stimuli (averaging over all trials and units) and b) applied a decoder to assess temporally evolving spike rate differences between conditions across and between individual trials, at the single unit level. The binwise neuronwise classification SVM decoder compared spike rate data from bins within single trial analysis windows containing different stimuli in the same context (both deviant), or the same stimuli in different contexts (standard vs deviant). Analysis windows were 1s wide, using bin widths of 50ms, based on a balance between temporal information and sufficient spike data in bins for classification accuracy (Fig S1). To determine if neurons significantly differentiated between stimuli in single trial spike rates, we statistically compared classifier accuracies to a shuffled control. Further, this single neuron decoding is not affected by spatial tonotopy and allows comparison of responses to different frequency stimuli in single neurons, given that we selected F1 & F2 stimuli such that both activate neurons equally, on average, across all simultaneously recorded neurons on a given electrode.

First, we found that A1 neurons encode contextual deviance during the oddball paradigm. We compared data containing the same tone but in different contexts (standard vs deviant; Fig 2D). We identified putative excitatory A1 neurons by regular spiking (RSN; Fig 2B, n=168) and identified mismatch-responsive RSNs (n=116). We found that A1 RSNs have larger responses to deviant stimuli than to standard stimuli for both different frequencies (F1 & F2), as expected, and that this was accompanied by significant differentiation of deviant and standard context for both F1 (Fig 2E left; t(115), p < 0.01, paired t-test) and F2 (Fig 2E center; t(115), p < 0.01, paired t-test). All layers of A1 differentiated stimuli based on context (Fig S2). Examining latencies, we observed the significance started at the 0-50ms bin after deviant onset and was long-lasting past the onsets of subsequent standard stimuli (Fig 2E, significance extending beyond 250 ms). Neurons in infragranular layers demonstrated a long-lasting modulation by context, from the deviant response to the next stimulus, and throughout much of the following train of standard stimuli (Fig S2). Context modulating single neuron responses and continuing into the subsequent stimulus may represent slow network dynamics as described previously for population encoding of frequency information (Shymkiv et al., 2025). The proposed function of this memory trace is to modulate responses to subsequent stimuli, and in agreement with this we also see altered rates and increased stimulus differentiation during the post-deviant standard.

Next, we found evidence that deviant responses *do not* convey a generalised surprise signal (i.e. no information of stimulus content). We compared ascending to descending deviant responses. Ascending and descending deviants evoked similar mean spike rate increases (Fig 2E right), yet our decoding analysis significantly differentiated between these conditions – i.e. A1 units differentiated between the two tones (F1 and F2) in the same context (Fig 2E right; t(115), p < 0.01, paired t-test). This result was confirmed with a linear mixed-effect model with Result/Shuffled as a fixed effect and Rat ID and Recording as a random effect (0-100ms window; ANOVA; F(1, 230) = 83.499, p < 0.001). Peak decoder accuracies of ascending vs descending deviants for individual neurons show a positively skewed distribution (Fig 2F). Significant encoding between ascending and descending deviants was confirmed in 39/116 neurons (Fig 2G).

We confirmed true deviance detection in our responses using the many-standards control. Deviant stimuli induced spike rate increases greater than those of many-standard stimuli and with significant decoding between the two (Fig 2I, t(115), p < 0.01, paired t-test). These data confirm that deviance detection is present (Fig 2I) and indicate that A1 RSNs responses are not only modulated by contextual deviance, but also other aspects (dev 1 vs dev 2; Fig 2E right). Thus, A1 RSNs likely encode some information of either the tone frequencies or frequency change between the ascending or descending oddball.

### Frequency selectivity cannot explain context encoding during the oddball paradigm

To resolve why deviants at F1 and F2 evoked different activity patterns (i.e. decoding in fig 2E) could be due to stimulus frequency encoding, we tested the hypothesis that A1 RSNs encode the small differences in frequency information during the oddball paradigm. We compared repeating standard stimuli of different frequencies (Fig 3A) and used data from the many-standards paradigm to compare stimuli which always had a certain tone at one timepoint (MS F1 & MS F2), to stimuli of random tone frequencies (RAND; Fig 3A). For MS F1 and F2, we centered analysis windows on with either the F1 or F2 frequencies, such that in all 40 trials the frequency of interest occurred at the same time within the analysis window. For the RAND condition, we centered analysis windows on the random stimuli preceding an F1 or F2 stimulus, which contain random frequency stimuli across the 40 trials. A1 RSNs did not differentiate between standard stimuli of F1 and F2 (Fig 3B left; t(115), p < 0.01, paired t-test), indicating no differentiation of stimuli based on frequency information. This may be because responses to standard stimuli were strongly attenuated (and nearly absent), likely due to strong stimulus specific adaptation, known to emerge upstream of A1. A1 RSNs also did not differentiate between F1 and F2 stimuli of the many-standards paradigm compared to random stimuli (Fig 3B center & right; t(115), p < 0.01, paired t-test). Notably, the two frequencies used (F1 and F2) were chosen for each neuron to be within the frequency response area, so individual A1 neurons not differentiating between these stimuli on a single trial basis is not unexpected because frequency encoding is predominantly achieved by spatial tonotopic arrangement. Interestingly, A1 responses to F1 and F2 were differentiable using these analyses only when they were contextually deviant (Fig 2E, right), which suggests either a) frequency encoding is sharpened when stimuli are contextually deviant or b) frequency is not encoded but, rather, deviations from context are encoded by A1 during the oddball paradigm. In summary, these data show A1 RSNs are not modulated by the small differences in stimulus frequency during the oddball paradigm when the stimulus was the repeated “standard”, and during the many standard control paradigm.

### A1 RS neurons encode the direction of frequency change

Next, we used a paradigm that allows us to assess how stimulus frequency change alters A1 responses. Cascade controls formed of sequentially increasing or decreasing frequency steps (Fig 3C) establish a regularity, as in the oddball, yet avoid repetition while maintaining a predictable context (Ruhnau et al., 2012). Comparing stimuli from the cascade paradigm centered on the same frequency allows determination of whether responses are modulated by frequency or frequency change direction, as throughout the analysis window there is a difference in paradigm context (ascending or descending stimulus set). When comparing ascending vs descending cascades, for stimuli other than center stimulus within the analysis window there is a difference in tone frequency (Fig 3D) – i.e for the first two stimuli in the analysis window, the tones in the descending cascade are higher frequency than those in the ascending cascade, and vice versa for the final two tones in the window. For the center tone in the analysis window, where the ascending and descending cascade cross at F1 or F2, we held the tone frequency equal. Thus, if A1 neurons differentiate between the center frequency in ascending or descending analysis windows, the neurons are responding to frequency change information, and not solely frequency (Fig 3D). In A1, mean spike rates were broadly similar for F1 or F2 when presented in ascending or descending cascades. However, neurons differentiated between the center frequency of these analysis windows, for both F1 and F2 as the center frequency (Fig 3E; t(115), p < 0.01, paired t-test). This result was confirmed by linear mixed-effect models with result/shuffled as a fixed effect and recording as a random effect, for both ascending (0-100ms window; ANOVA; F(1, 230) = 19.736, p < 0.001) and descending (0-100ms window; ANOVA; F(1, 230) = 16.017, p < 0.001). Different response patterns evoked by ascending and descending cascades were confirmed in 16/116 neurons (13.8%) for F1 and 13/116 neurons (11.2%) for F2 (Fig 3F-G). A1 RSNs exhibit sensitivity to cascade direction at the center tone, consistent with modulation by recent frequency-change. To further assess how recent frequency change information may be encoded we used a constrained many-standards paradigm where the frequency change preceding F1 or F2 is equal in the control and oddball conditions (i.e., when F2 in the control was preceded by F1, or vice versa; Fig S3). Responses to the oddball deviant were larger than constrained many-standards responses and decoding analyses evinced a distinct response across neurons, indicating that A1 is not solely responding to short-duration frequency changes, but also the context of the current paradigm.

### Deviance detection errors as a difference signal

The above results demonstrate that A1 responses to deviant stimuli encode contextual deviance, rather than merely the tone frequency (Figs 1; Fig 3) or its difference from the previous stimulus (Fig 3; Fig S4). We next tested if augmented deviant responses act as a difference signal (as in the predictive coding framework (Rao and Ballard, 1999; Friston, 2005; Clark, 2013)) or enhanced sensory response (as observed in other paradigms (Muzzu and Saleem, 2021; Furutachi et al., 2024)). We reasoned that if deviance detection reflects purely an enhanced sensory response, then deviant responses to a given frequency would be equal even when varying the frequency of the standard. Conversely, if the deviant response is different when there is a larger frequency jump from the standard, then this would suggest a difference signal – i.e. that the augmented response reflects a difference between what is predicted and what is presented. In other words, if one holds the deviant stimulus features constant, but varies the standard stimulus features (i.e. the “prediction”), is the response to the deviant different? We recorded A1 neurons (n’s = 810 total; 156 RSNs; 75 mismatch-responsive RSNs) from 21 electrode positions; 1344 total channels) from 2 female rats. We presented a fixed-frequency deviant but varied the magnitude of frequency difference between the deviant and standard stimuli (Fig 4A). In this oddball we the standard sequence theoretically establishes an internal prediction centered on the standard frequency and standard/deviant step size is a proxy for prediction/input discrepancy. A1 mismatch-responsive RSNs significantly differentiated between deviants in runs of varied standards, both for differences in different directions (Fig 4B; t(74), p < 0.01, paired t-test) and opposite directions (Fig 4C). We also found larger mean spike rate increases to ascending 1 octave than 0.5 octave stimuli, though this effect was not present in the descending condition (Fig 4D; t(74) p < 0.01, paired t-tests). Comparing data from the ascending or descending 1 octave conditions, we found a moderate positive correlation between mean deviant responses (Fig 4E-F). These results indicate that some neurons have larger deviance detection, but the deviance detection is not equal across conditions tested. Data for all comparisons are available in Fig S5.

Somatosensory studies further show that the timing of first spikes in primary sensory neurons convey rich information about the stimulus (Petersen et al., 2001; Johansson and Birznieks, 2004). We reasoned that if prediction-error responses similarly contained information about stimulus characteristics (in this case, the magnitude of deviant frequency change), this may be reflected by first spike latency (FSL), with earlier FSL corresponding to a larger frequency difference. Conversely, if prediction-errors encoded sensory enhancement or invariant surprise, we would expect no differences between deviant responses. We compared FSLs (Fig 4G-H) and a two-way repeated-measures ANOVA with factors frequency change Magnitude (full, half) and change Direction (ASC vs DES) showed a significant, non-crossover Magnitude × Direction interaction. The marginal means for the effect of direction were not different (p=0.13) but were for the effect of change magnitude (p=0.0003). Follow-up paired comparisons showed that response latencies were significantly faster for full than half octaves for both the ascending (t(29)=−3.23, p=0.003), and descending deviants (t(29)=−2.19, p=0.037).

Given that the deviant was always a fixed frequency for each neuron, here a population decoder could be used without the confound of decoding tonotopic frequency information. We used a trial-wise Maximum correlation classifier, and revealed the sampled neural population differentiates between ascending and descending 1 octave deviant responses (Fig 4), indicating that despite the constant deviant tone, population response patterns are varied when the STD is altered. To confirm that deviant responses contain information beyond a residual frequency information from the previous stim, we also tested for differentiation between Direction (ASC vs DES) or Magnitude (0.5 vs 1 oct) for each trial at different time windows (STD, post-STD and DEV) (Fig 4J). While decoder accuracy of combined Magnitude and Direction for STD and post-STD time windows was above chance, there was a large increase in decoder accuracy in the DEV time window, confirming a deviant-specific difference signal. In summary, these data demonstrate that A1 deviance detection responses vary with the difference between internal prediction and sensory input, results consistent with a difference calculation under the predictive coding framework.

## DISCUSSION

Augmented neural responses to deviant stimuli have been extensively characterised (Dürschmid et al., 2016; Parras et al., 2017; Hockley et al., 2025; Shymkiv et al., 2025), but here we go beyond response magnitude to investigate the information contained by these neural responses. We compared A1 neural responses to auditory oddball stimuli differing in either frequency or context, or a combination of both. Using SVM decoding of single-trial spike data, we identified differences in activity patterns of individual neurons that are not present when comparing mean spike rates. Importantly, this approach demonstrated that A1 RSNs encode contextual deviance as a prediction-input difference during the oddball paradigm, not simply stimulus features or a generalised surprise signal. These results have implications for how signals observed during oddball sequences – a ubiquitous experimental paradigm used in basic and clinical neuroscience – should be interpreted.

Cortical pyramidal neuron deviance detection could represent an invariant surprise signal, an enhanced sensory input (Muzzu and Saleem, 2021; Furutachi et al., 2024) or the difference between the internal prediction and the unexpected, incoming sensory input (Rao and Ballard, 1999; Friston, 2005; Clark, 2013). Here we designed a series of experimental paradigms to systematically distinguish between these possibilities. We show that A1 RSNs differentiate between ascending and descending deviants, indicating that deviance detection contains some stimulus information and therefore is not an invariant surprise signal. Using a fixed deviant oddball, we found an enhanced deviance detection with increased standard-deviant frequency difference, which could be analytically decoded from single neurons or the population response. Because the deviant stimulus remained physically identical across conditions, this result confirms that deviance detection is not a simple enhancement of sensory information. Instead, our findings support the interpretation that neural deviance detection conveys information about the difference between internal prediction and sensory input, consistent with deviance detection being interpreted as a prediction error in a predictive coding framework. The neural mechanisms underlying this computation remain unknown. One possibility is that enhancement of frequency change information contributes to sensory prediction errors. Such a mechanism would account for neuronal responses encoding both the direction of frequency change (ascending vs descending deviants; Figs 1 & 3) in addition to the magnitude of frequency change (Fig 4). While increased deviant responses with increased standard-deviant frequency spacing have been reported previously (Ulanovsky et al., 2003), those findings were obtained using a classical oddball paradigm. In contrast, our fixed-deviant oddball paradigm is better controlled for studying difference signals, as the tested deviant frequency remains constant, not varied with larger standard-deviant frequency spacing, thereby manipulating the standard–deviant frequency difference without changing the deviant stimulus itself.

Recent evidence using Ca^2+^ imaging during an attended VR sensory-spatial paradigm posits prediction errors as enhanced sensory responses (Furutachi et al., 2024). To determine whether differences in the temporal resolution of spike data and Ca²⁺ imaging could explain the discrepancy between our results and those of Furutachi et al (Furutachi et al., 2024), we repeated an analysis with degraded temporal information (Fig S6). Under these conditions, neural deviance detection nevertheless conveyed a difference signal. Accurate classification with less accurate temporal information suggests that temporal resolution is unlikely to explain the difference between the present results and those collected with 2-photon imaging. Therefore, we consider differences in the paradigms used to be the most likely explanation, with the simple unattended sensory prediction errors involving lower-order processing than the spatial-sensory prediction errors used by Furutachi et al (Furutachi et al., 2024).

Interestingly, A1 neuronal populations encode sensory information after a stimulus terminates, i.e. beyond stimulus offset, extending into the subsequent stimulus (Shymkiv et al., 2025). This has been interpreted as a network hysteresis, or a memory trace of the past stimulus information persisting in population activity. Such “slow network dynamics” are proposed to modulate responses to subsequent stimuli, and as such may underlie local transitional responses (Maheu et al., 2019; Meyniel et al., 2016; Walker et al., 2023). Cortical slow encoding of contextual deviance may therefore be a key mechanism of predictive coding – at least in certain cases, such as the “oddball” paradigm. This contextual deviance is encoded in a slow trace, extending beyond the end of the stimulus and into the subsequent stimulus, as we found in RSNs. We did not find evidence for a slow memory trace in FSNs (Fig S4), suggesting that this effect is specific to RSNs, consistent with slow dynamics of pyramidal neurons.

We found that the small frequency differences in our oddball paradigm were not well differentiated well by single A1 RSNs. This may be explained by the fact that in the stimuli used here, F1 and F2 were both within the receptive fields of the neurons, with half an octave frequency spacing, meaning that both stimuli could activate any given neuron. This result supports the conclusion that the small frequency differences during the classical oddball paradigm did not underlie what we interpret as context-dependent modulation of neural responses. The bin size used for analysis (50 ms) is too large to detect precise spike timing, but the auditory cortex relies on a rate code (Buck et al., 2025) and frequency is encoded by rates in A1 (Micheyl et al., 2013). Therefore, the bin widths would not be expected to greatly affect the detection of frequency encoding presented here. While smaller bins may have some advantages, they would also reduce the number of spikes per bin and require more stimulus presentations to maintain the decoder accuracy.

A1 inhibitory interneurons also respond to auditory deviants (Pérez-González et al., 2021; Yarden et al., 2022) and they contribute to deviance detection by bidirectionally modulating pyramidal neuron responses to deviants (Hamm and Yuste, 2016; Hertäg and Sprekeler, 2020; Yarden et al., 2022; Bastos et al., 2023; Furutachi et al., 2024). Here, while we cannot distinguish between interneuron cell types, we can study the information content separately in putative pyramidal and certain types of inhibitory interneurons identified by their spike waveforms. As with RSNs, FSNs show deviance detection (Fig S4D; deviant vs many-standards). We show that FSNs, like the RSNs, also differentiate between ascending and descending deviants, suggesting that deviance detection in this subpopulation is also unlikely to reflect an invariable surprise signal.

While our present study focused on A1, other downstream brain regions are also of great interest. Auditory oddball prediction errors are also present in secondary auditory areas (Nieto-Diego and Malmierca, 2016), and the circuit connecting the anterior cingulate cortex to the primary sensory cortex contributes to the generation of prediction errors in the primary sensory cortex (Bastos et al., 2023; Hockley and Malmierca, 2024; Hockley et al., 2025; Peterka et al., 2026). While we reveal here that deviance detection represents a prediction-based difference signal in A1 RSNs, this difference information may result in enhancement of sensory information in downstream (or even upstream) regions and circuits, which could concord with the study by (Furutachi et al., 2024). It may also be possible that enhancement of sensory response may occur in some neurons in proportion to the standard/deviant difference. While we found no evidence for this here, such a mechanism could underlie enhanced responses to deviants in higher-order areas, reflecting information about the new stimulus rather than signalling a prediction error.

Our study has limitations that merit mention. Here, we recorded under anesthesia, which can alter aspects of prediction error processing (Quaedflieg et al., 2014; Nourski et al., 2018; Xiong et al., 2024; Bohórquez et al., 2025), although urethane maintains balanced inhibitory/excitatory neuronal activity (Hara and Harris, 2002) and has a robust history of simple sensory prediction error research (Nieto-Diego and Malmierca, 2016; Parras et al., 2017; Hockley et al., 2025). Nevertheless, future studies on the information content of prediction errors in awake animals are warranted. That said, the influence of selective attention on sensory response gain is well established, and an anesthetised preparation such as ours may mitigate such influences. This study also focused on A1, so further studies of non-lemniscal auditory areas would be of interest. The data used here also come solely from female animals; future work should extend recordings to male rats, although our past work has not found sex differences in adult cortical deviance detection (e.g. (Rader Groves et al., 2026). One advantage of the present analysis is that it classifies individual trials, so significant decoding indicates that contextual deviance is contained within single bin data from single stimulus trials. This results in moderate decoder accuracy, yet significantly above chance. Decoding accuracy could likely be further improved by increasing the number of stimulus presentations per neuron, thereby providing more data for training the classifier.

In summary, we show that A1 RSN deviance detection contains information about frequency change, which is expressed as a difference signal in a fixed-deviant oddball paradigm. A1 deviance detection scales with the difference between internal predictions and unexpected sensory inputs, therefore providing direct evidence that deviance detection reflects cortical sensory prediction errors, as defined by predictive coding frameworks. Revealing that neural deviance detection contains a fundamental hallmark of predictive coding frameworks strengthens the interpretation of deviance detection as a sensory prediction error. Deviance detection provides a valuable tool for investigating low-level predictive coding, as basic sensory deviations elicit low-latency sensory prediction errors.

## Acknowledgments

This project has received funding from the European Union’s Horizon 2020 research and innovation program under the Marie Skłodowska-Curie grant agreement No PROOPI340-USAL4EXCELLENCE to AH and MSM; PID2023-148541OB-I00 funded by MICIU/AEI and FEDER EU to MSM; SA218P23 Funded by Consejería de Educación, Junta de Castilla y León; and CLU-2023-1-01 funded by the Regional Government of Castile and León & the ERDF Operational Programme to MSM. NIH R01EY033950 to JPH; The authors would like to thank Laura H Bohórquez for surgical and histological assistance and comments on an earlier version of this manuscript.

## Author contributions

Designed research, AH; Performed research, AH; Analyzed data, AH and CGG; Wrote the first draft of the paper, AH; Edited the paper, AH, CGG, JPH and MSM.

## SUPPLEMENTAL INFORMATION

**Figure S1:**
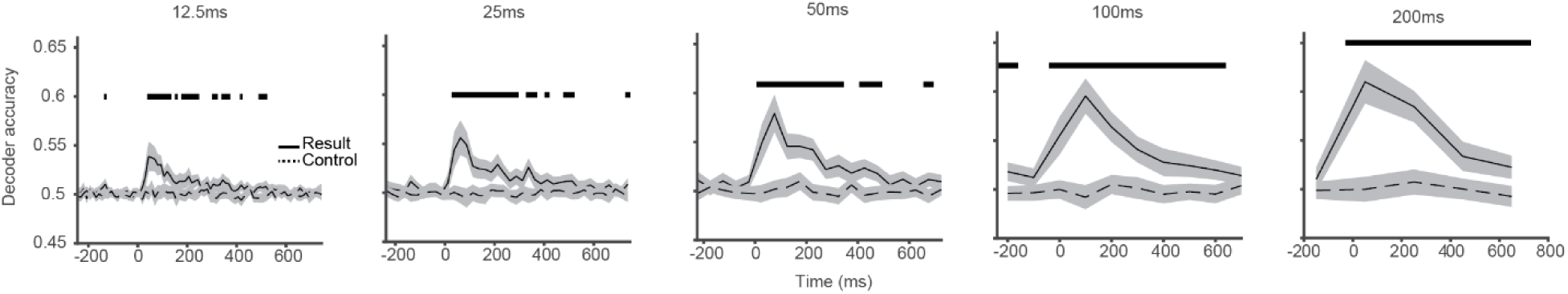
Effect of bin size on decoding scores. Decoder for the comparison in Fig 2E right (descending vs ascending deviant) was run with different spike bin windows.

**Figure S2:**
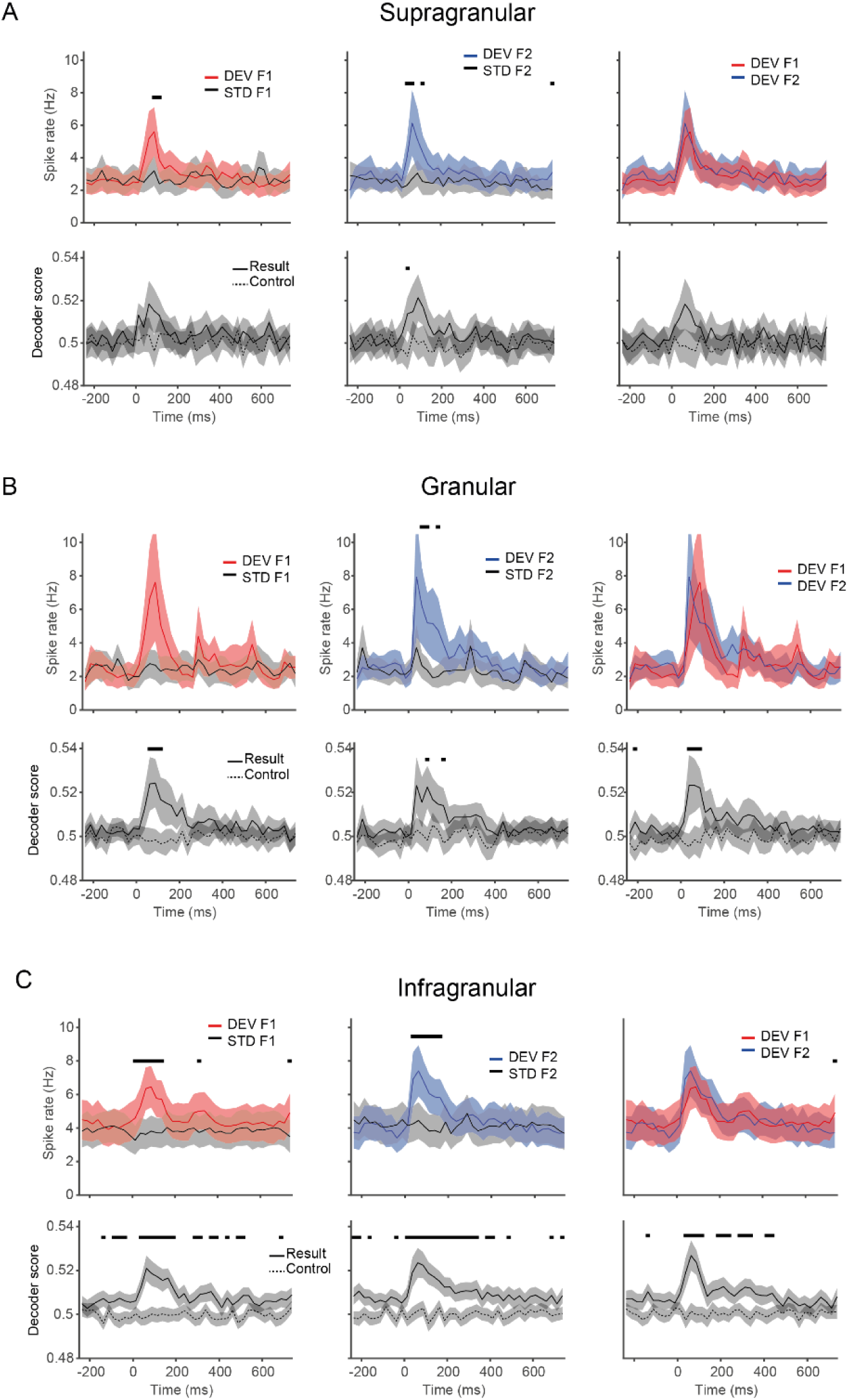
Layer-specific encoding of context in A1. Using the depth of neurons on the electrode shank, all neurons were separated by putative cortical layers of supragranular (A), granular (B), and infragranular (C). n’s = 112, 133 & 392, respectively. Shaded areas represent 95% CI. Black bars indicate p < 0.01, binwise Bonferroni-corrected paired t-test. Analysis as in Fig 2E reveals our A1 result is largely due to infragranular layer neurons, and to a lesser degree, granular layer neurons.

**Fig S3:**
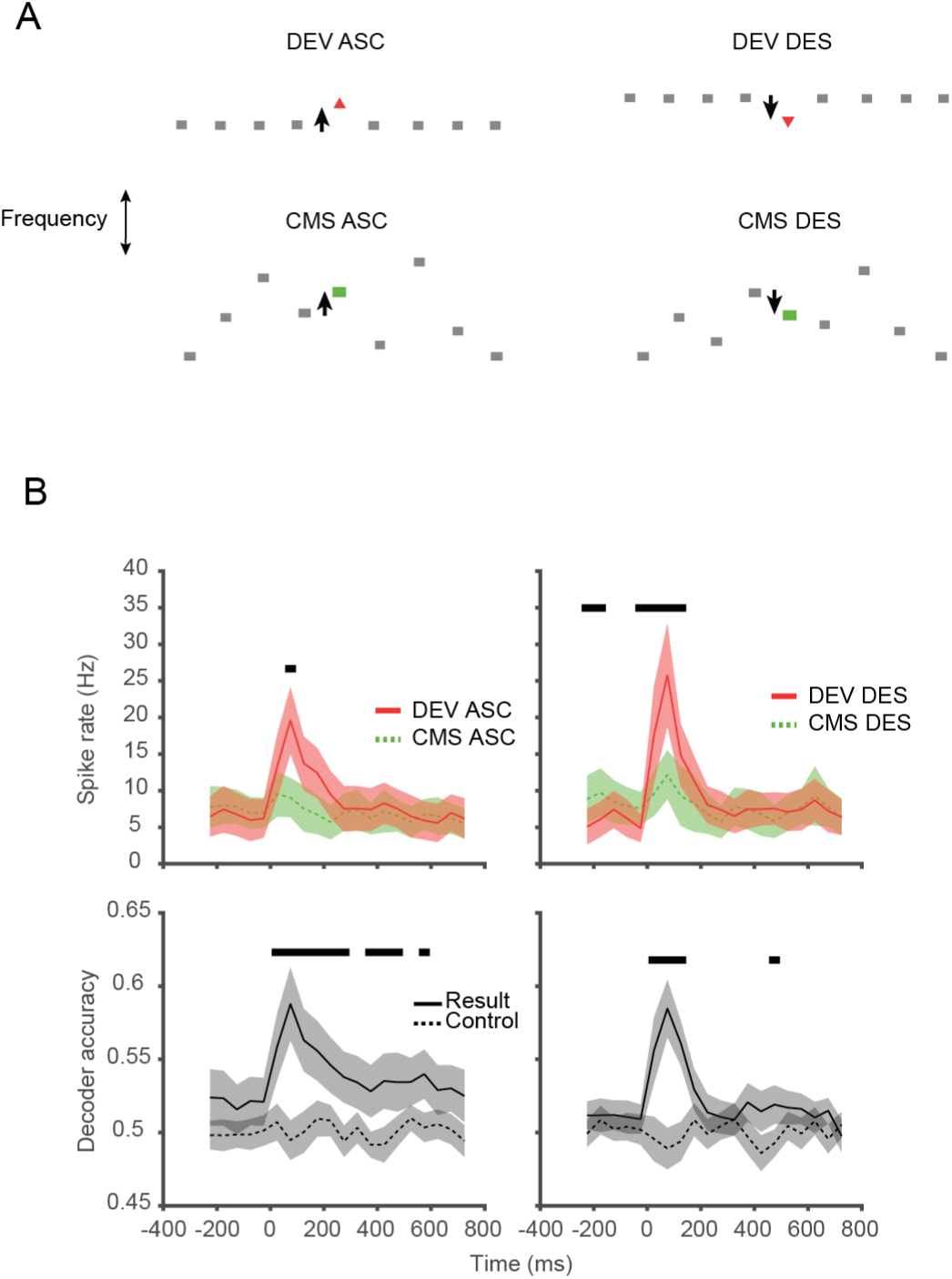
A) The constrained many-standards paradigm with equal frequency changes preceding frequencies of interest. B) For both ascending and descending conditions, deviant responses are larger than CMC responses, and with significant decoding between the two. Shaded areas represent 95% CI. Black bars indicate p < 0.01, paired t-test; n = 72 mismatch-responsive RSNs.

**Figure S4:**
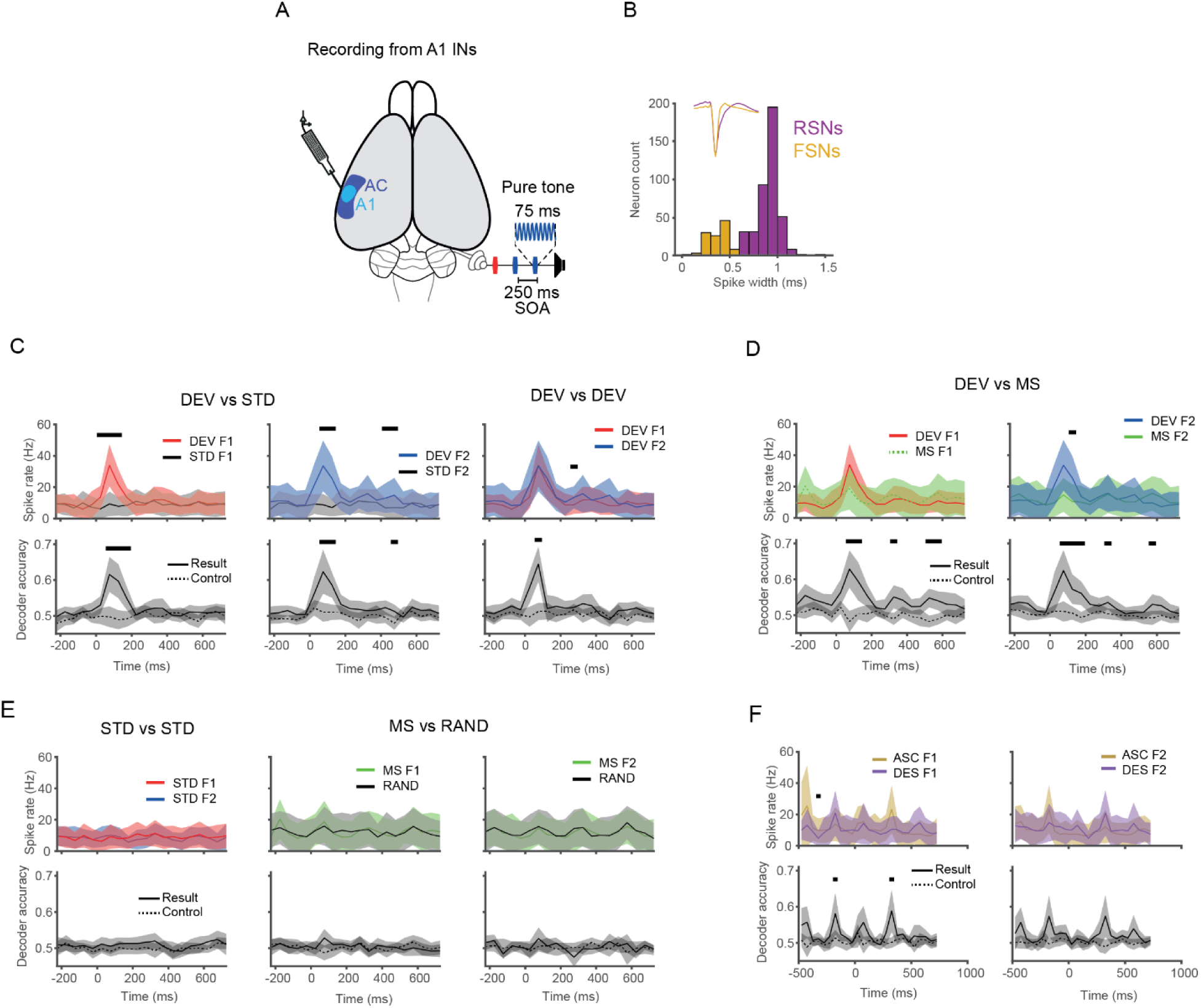
FSN responses encode deviance detection but no memory trace. A) Schematic of multichannel recording locations in A1. B) Neurons were separated by spike width (positive to negative peak duration; cutoff 0.6ms) into regular spiking (RSNs; putative excitatory) and fast spiking (FSNs; putative inhibitory) neurons. C) Comparison of deviant to standard responses (mismatch) D) Comparison of deviant to man-standard responses (deviance detection) E) Comparison of same context but different frequency information. F) Comparison of ascending to descending cascades (frequency change information). Shaded areas represent 95% CI. Black bars indicate p < 0.01, binwise paired t-test. n = 24 mismatch-responsive FSNs.

**Figure S5:**
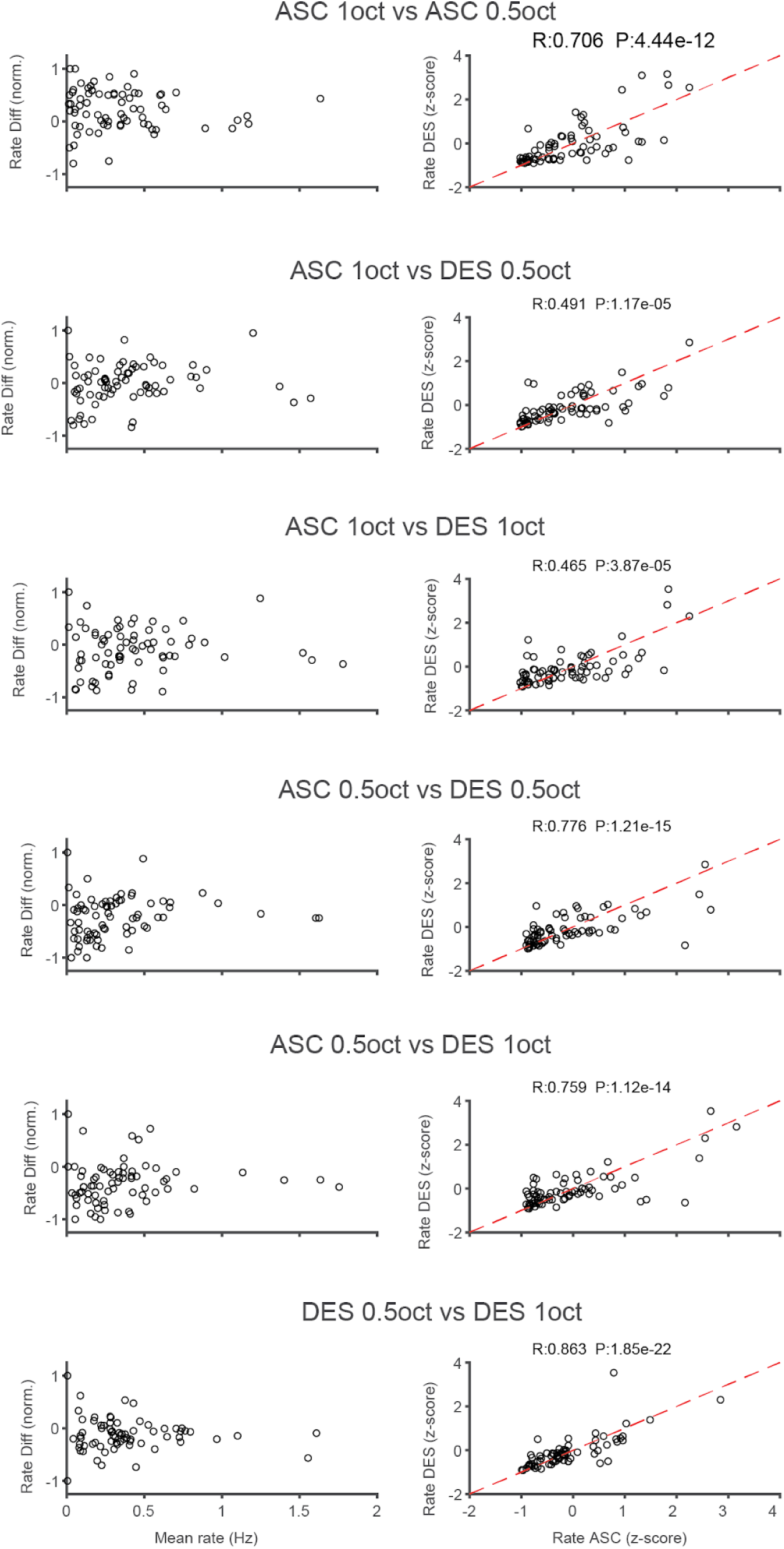
A1 difference signals. For all comparisons, correlation and Bland-Altman plots show neurons with greater deviant responses are more likely to have larger firing rates between fixed deviants during variable standards. Dots represent individual neurons; n = 72 mismatch-responsive RSNs.

**Figure S6:**
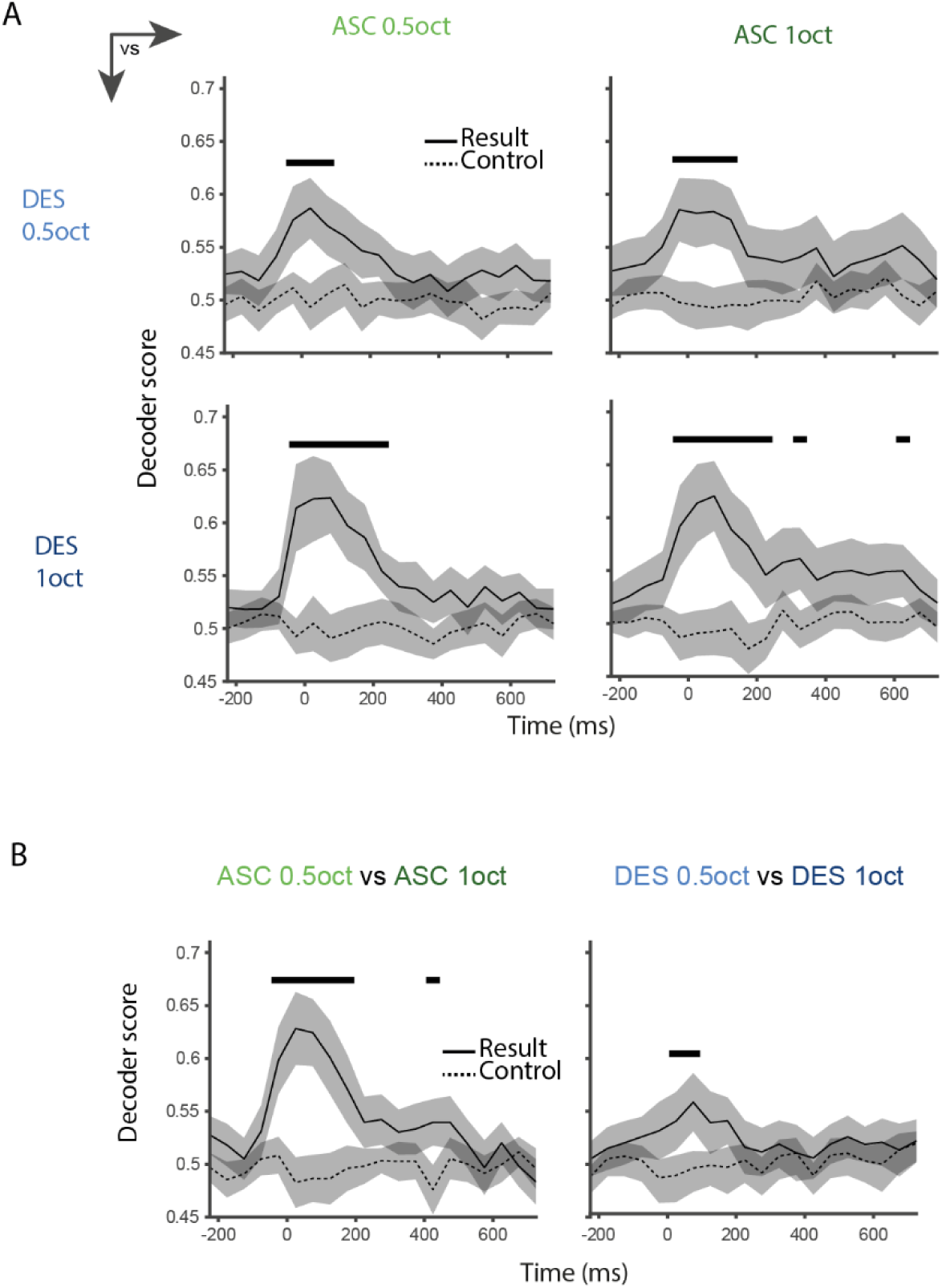
Mimicking Ca^2+^ imaging dynamics. To determine if temporal acuity contributed to our results differing from those of Furutachi et al, we ran the same analyses as Fig 4B,C but temporal information was degraded using 50ms bin widths then smoothed with a gaussian-weighted average over 5 bins. Shaded areas represent 95% CI. Black bars indicate p < 0.01, paired t-test; n = 72 mismatch-responsive RSNs.

## Notes

Conflict of interest statement:The authors declare no competing financial interests.

### Competing Interest Statement

The authors have declared no competing interest.

### Summary of Updates

Updated manuscript with new analyses and writing throughout

## REFERENCES

Aizenbud I et al. (2025) Neural mechanisms of predictive processing: a collaborative community experiment through the OpenScope program. arXiv [q-bioNC] Available at: 10.48550/arXiv.2504.09614.

Arnal LH, Giraud AL (2012) Cortical oscillations and sensory predictions. Trends Cogn Sci 16:390–398.

Asari H, Zador AM (2009) Long-lasting context dependence constrains neural encoding models in rodent auditory cortex. J Neurophysiol 102:2638–2656.

Asko O, Blenkmann AO, Leske SL, Foldal MD, Llorens A, Funderud I, Meling TR, Knight RT, Endestad T, Solbakk AK (2024) Altered hierarchical auditory predictive processing after lesions to the orbitofrontal cortex. Elife 13:1–29.

Audette NJ, Zhou WX, Chioma AL, Schneider DM (2022) Precise movement-based predictions in the mouse auditory cortex. Curr Biol 32:4925–4940.e6.

Bastos AM, Usrey WM, Adams RA, Mangun GR, Fries P, Friston KJ (2012) Canonical microcircuits for predictive coding. Neuron 76:695–711.

Bastos G, Holmes JT, Ross JM, Rader AM, Gallimore CG, Wargo JA, Peterka DS, Hamm JP (2023) Top-down input modulates visual context processing through an interneuron-specific circuit. Cell Rep 42:113133.

Bohórquez LH, Malmierca MS, Hockley A (2025) Do auditory deviants evoke cortical state changes under anaesthesia? A proof-of-concept study. Exp Physiol Available at: 10.1113/EP093378.

Buccino AP, Hurwitz CL, Garcia S, Magland J, Siegle JH, Hurwitz R, Hennig MH (2020) Spikeinterface, a unified framework for spike sorting. Elife 9:1–24.

Buck A, Dupont T, Cavanagh RA, Postal O, Bourien J, Puel JL, Michalski N, Gourévitch B (2025) Neural Response Reliability as a Marker of the Transition of Neural Codes along Auditory Pathways. Adv Sci (Weinh) Available at: 10.1002/advs.202508777.

Clark A (2013) Whatever next? Predictive brains, situated agents, and the future of cognitive science. Behav Brain Sci 36:181–204.

Dürschmid S, Edwards E, Reichert C, Dewar C, Hinrichs H, Heinze HJ, Kirsch HE, Dalal SS, Deouell LY, Knight RT (2016) Hierarchy of prediction errors for auditory events in human temporal and frontal cortex. Proc Natl Acad Sci U S A 113:6755–6760.

Englitz B, Akram S, Elhilali M, Shamma S (2024) Decoding contextual influences on auditory perception from primary auditory cortex. Elife 94296 Available at: https://app.dimensions.ai/details/publication/pub.1167336503.

Espejo ML, David SV (2024) A sparse code for natural sound context in auditory cortex. Curr Res Neurobiol 6 Available at: 10.1016/j.crneur.2023.100118.

Fiáth R, Márton AL, Mátyás F, Pinke D, Márton G, Tóth K, Ulbert I (2019) Slow insertion of silicon probes improves the quality of acute neuronal recordings. Sci Rep 9:1–17.

Friston K (2005) A theory of cortical responses. Phil Trans R Soc B 360:815–836.

Furutachi S, Franklin AD, Aldea AM, Mrsic-Flogel TD, Hofer SB (2024) Cooperative thalamocortical circuit mechanism for sensory prediction errors. Nature 633:398–406.

Furutachi S, Hofer SB (2026) Rethinking predictive processing. Annu Rev Neurosci Available at: 10.1146/annurev-neuro-102124-031410.

Gallimore CG, Ricci DA, Hamm JP (2023) Spatiotemporal dynamics across visual cortical laminae support a predictive coding framework for interpreting mismatch responses. Cereb Cortex 33:9417–9428.

Gelens F, Äijälä J, Komatsu M, Uran C, Jensen MA, Miller KJ, Ince RAA, Vinck M, Canales-Johnson A (2024) Distributed representations of prediction error signals across the cortical hierarchy are synergistic. Nat Commun 2024 Available at: 10.1038/s41467-024-48329-7.

Gershman SJ, Assad JA, Datta SR, Linderman SW, Sabatini BL, Uchida N, Wilbrecht L (2024) Explaining dopamine through prediction errors and beyond. Nat Neurosci 27:1645–1655.

Hamm JP, Shymkiv Y, Han S, Yang W, Yuste R (2021) Cortical ensembles selective for context. Proc Natl Acad Sci U S A 118:1–12.

Hamm JP, Yuste R (2016) Somatostatin Interneurons Control a Key Component of Mismatch Negativity in Mouse Visual Cortex. Cell Rep 16:597–604.

Han S, Helmchen F (2024) Behavior-relevant top-down cross-modal predictions in mouse neocortex. Nat Neurosci 27:298–308.

Hara K, Harris RA (2002) The anesthetic mechanism of urethane: The effects on neurotransmitter-gated ion channels. Anesth Analg 94:313–318.

Harms L, Fulham WR, Todd J, Budd TW, Hunter M, Meehan C, Penttonen M, Schall U, Zavitsanou K, Hodgson DM, Michie PT (2014) Mismatch negativity (MMN) in freely-moving rats with several experimental controls. PLoS One 9 Available at: 10.1371/journal.pone.0110892.

Hertäg L, Sprekeler H (2020) Learning prediction error neurons in a canonical interneuron circuit. Elife 9 Available at: 10.7554/eLife.57541.

Hockley A, Bohórquez LH, Malmierca MS (2025) Top-down prediction signals from the medial prefrontal cortex govern auditory cortex prediction errors. Cell Rep 44 Available at: 10.1016/j.celrep.2025.115538.

Hockley A, Malmierca MS (2024) Auditory processing control by the medial prefrontal cortex: A review of the rodent functional organisation. Hear Res 443:108954.

Johansson RS, Birznieks I (2004) First spikes in ensembles of human tactile afferents code complex spatial fingertip events. Nat Neurosci 7:170–177.

Keller GB, Mrsic-Flogel TD (2018) Predictive Processing: A Canonical Cortical Computation. Neuron 100:424–435.

Maheu M, Dehaene S, Meyniel F (2 2019) Brain signatures of a multiscale process of sequence learning in humans. Elife 8 Available at: 10.7554/ELIFE.41541.

Meyniel F, Maheu M, Dehaene S (12 2016) Human Inferences about Sequences: A Minimal Transition Probability Model. PLoS Comput Biol 12:e1005260.

Micheyl C, Schrater PR, Oxenham AJ (2013) Auditory Frequency and Intensity Discrimination Explained Using a Cortical Population Rate Code. PLoS Comput Biol 9 Available at: 10.1371/journal.pcbi.1003336.

Muzzu T, Saleem AB (2021) Feature selectivity can explain mismatch signals in mouse visual cortex. Cell Rep 37:109772.

Nieto-Diego J, Malmierca MS (2016) Topographic Distribution of Stimulus-Specific Adaptation across Auditory Cortical Fields in the Anesthetized Rat. PLoS Biol 14:1–30.

Nourski KV, Steinschneider M, Rhone AE, Kawasaki H, Howard MA, Banks MI (2018) Auditory predictive coding across awareness states under anesthesia: An intracranial electrophysiology study. J Neurosci 38:8441–8452.

Pachitariu M, Sridhar S, Pennington J, Stringer C (2024) Spike sorting with Kilosort4. Nat Methods 21:914–921.

Parras GG, Nieto-Diego J, Carbajal GV, Valdés-Baizabal C, Escera C, Malmierca MS (2017) Neurons along the auditory pathway exhibit a hierarchical organization of prediction error. Nat Commun 8 Available at: 10.1038/s41467-017-02038-6.

Pérez-González D, Parras GG, Morado-Díaz CJ, Aedo-Sánchez C, Carbajal GV, Malmierca MS (2021) Deviance detection in physiologically identified cell types in the rat auditory cortex. Hear Res 399:107997.

Peterka DS, Imai F, Ross JM, Bastos G, Hornick MA, Rachmany L, Gallimore CG, Hockley A, Hamm JP (2026) Anterior cingulate feedback facilitates rapid global context modulation in primary visual cortex. Cell Rep 45:118151.

Petersen RS, Panzeri S, Diamond ME (2001) Population coding of stimulus location in rat somatosensory cortex. Neuron 32:503–514.

Quaedflieg CW, Münte S, Kalso E, Sambeth A (2014) Effects of remifentanil on processing of auditory stimuli: A combined MEG/EEG study. J Psychopharmacol 28:39–48.

Rader Groves AM, Ricci DA, Wargo JA, Florisel VI, Dalwai HS, Ferrell AD, Desai B, Ross JM, Gallimore CG, West CL, Bastos G, Bolton JL, Imai F, Hamm JP (2026) Mismatch negativity matures in adolescence and without sensitivity to microglial depletion. Neuropsychopharmacology Available at: 10.1038/s41386-026-02547-3.

Rao RPN, Ballard DH (1999) Predictive coding in the visual cortex: A functional interpretation of some extra-classical receptive-field effects. Nat Neurosci 2:79–87.

Ruhnau P, Herrmann B, Schröger E (2012) Finding the right control: The mismatch negativity under investigation. Clin Neurophysiol 123:507–512.

Schröger E, Wolff C (1996) Mismatch response of the human brain to changes in sound location. Neuroreport 7:3005–3008.

Schultz W (2016) Dopamine reward prediction error coding. Osaka J Math 18:23–32.

Shymkiv Y, Hamm JP, Escola S, Yuste R (2025) Slow cortical dynamics generate context processing and novelty detection. Neuron:1–11.

Smith PH, Uhlrich DJ, Manning KA, Banks MI (2012) Thalamocortical projections to rat auditory cortex from the ventral and dorsal divisions of the medial geniculate nucleus. J Comp Neurol 520:34–51.

Ulanovsky N, Las L, Nelken I (4 2003) Processing of low-probability sounds by cortical neurons. Nat Neurosci 6:391–398.

Walker EY, Pohl S, Denison RN, Barack DL, Lee J, Block N, Ma WJ, Meyniel F (2023) Studying the neural representations of uncertainty. Nat Neurosci 26:1857–1867.

Watabe-Uchida M, Eshel N, Uchida N (2017) Neural circuitry of reward prediction error. Annu Rev Neurosci 40:373–394.

Xiong Y (sophy), Donoghue JA, Lundqvist M, Mahnke M, Major AJ, Brown EN, Miller EK, Bastos AM (2024) Propofol-mediated loss of consciousness disrupts predictive routing and local field phase modulation of neural activity. Proceedings of the National Academy of Sciences 121 Available at: 10.1073/pnas.

Yarden TS, Mizrahi A, Nelken I (2022) Context-Dependent Inhibitory Control of Stimulus-Specific Adaptation. Journal of Neuroscience 42:4629–4651.

